# Convergent DNA Methylation Abnormalities at Bivalent Chromatin in Human Growth Disorders

**DOI:** 10.1101/2025.07.08.663614

**Authors:** Marie E. Strauss, Yoshiko Takahashi, Jihye Lee, Camille Perez, Xiaoting Chen, Yuri Lee, Zachary S. Pope, Hongseok Yun, Matthew T. Weirauch, Minji Byun

## Abstract

Loss-of-function mutations in *DNMT3A*, a DNA methyltransferase, or *NSD1*, a histone methyltransferase, cause overgrowth syndromes. Conversely, disruption of the DNMT3A domain that binds NSD1-deposited H3K36 dimethylation (H3K36me2) results in growth restriction. To investigate the molecular basis of these opposing growth outcomes, we generated isogenic human embryonic stem cells carrying growth syndrome–associated mutations in *DNMT3A* and *NSD1*. Unexpectedly, both overgrowth- and growth restriction– associated *DNMT3A* mutations led to DNA hypomethylation in a shared subset of active enhancers, implicating H3K36me2 in directing enhancer methylation maintenance. In contrast, bivalent chromatin—marked by both active and repressive chromatin modifications—showed divergent DNA methylation changes: hypermethylation in growth restriction-associated *DNMT3A* mutants and hypomethylation in overgrowth-associated *DNMT3A* or *NSD1* loss-of-function mutants. These findings identify locus-specific DNA methylation defects as a common molecular feature and nominate dysregulated DNA methylation at bivalent chromatin as a potential driver of abnormal growth phenotypes.

## Introduction

*DNMT3A* encodes a de novo DNA methyltransferase essential for establishing DNA methylation patterns during development [1]. Germline mutations in *DNMT3A* underlie two distinct Mendelian syndromes with opposing growth phenotypes. Heterozygous loss-of-function (LoF) mutations cause Tatton-Brown-Rahman Syndrome (TBRS; OMIM 615879), characterized by tall stature, macrocephaly, intellectual disability, and dysmorphic features [2], [3]. Conversely, specific heterozygous missense mutations in the PWWP (Pro-Trp-Trp-Pro) domain have been linked to Heyn-Sproul-Jackson syndrome (HESJAS; OMIM 618724), marked by growth restriction, microcephaly, and impaired intellectual development [4], [5]. The PWWP domain recognizes di- or tri-methylated lysine 36 of histone H3 (H3K36me2/3) [6], [7], while the isoform-specific N-terminal region of DNMT3A1 binds monoubiquitinated lysine 119 of histone H2A (H2AK119ub1), directing genomic targeting [8], [9]. The catalytic methyltransferase domain, located near the C-terminus, is autoinhibited by the ADD (ATRX-Dnmt3-Dnmt3L) domain. When the ADD domain binds unmodified H3K4, it relieves autoinhibition and enables catalytic activity [10]. Together, these domains coordinate the recruitment and enzymatic activity of DNMT3A in a chromatin-context-dependent manner.

Intriguingly, mutations in other epigenetic regulators produce growth disorders with clinical features overlapping those of *DNMT3A*-related syndromes [11]. For example, LoF mutations in *NSD1*, a histone methyltransferase that deposits H3K36me1/2 [12], cause Sotos syndrome (OMIM 117550), the most common overgrowth-intellectual disability syndrome [11], [13], [14], [15], whereas *NSD1* copy number gains due to microduplications are associated with microcephaly, short stature and developmental delay [16], [17]. Similarly, LoF mutations in *EZH2*, *EED*, *SUZ12*–each encoding core components of the Polycomb Repressive Complex (PRC) 2 that deposit H3K27me3–cause overgrowth-intellectual disability syndromes: Weaver syndrome (OMIM 277590), Cohen-Gibson syndrome (OMIM 617561), and Imagawa-Matsumoto syndrome (OMIM 618786), respectively [18], [19], [20], [21], [22].

Genome-wide DNA methylation studies have identified distinct DNA methylation abnormalities associated with the *DNMT3A*-related growth syndromes. Samples from overgrowth TBRS patients display focal hypomethylation at enhancers, Polycomb repressed regions, and developmental genes such as homeobox domains [23], [24], [25]. In contrast, samples from growth restriction HESJAS patients show hypermethylation at Polycomb-marked DNA methylation valleys and key developmental genes [4]. Despite these insights, differences in cell type, age, sex, and genetic background in patient-derived samples complicate direct comparisons across syndromes. Moreover, because these studies analyzed fully differentiated tissues such as blood, it remains unclear when during development the DNA methylation defects arise.

To address these limitations, we introduced TBRS- and HESJAS-associated *DNMT3A* mutations into human embryonic stem cells (hESCs), enabling assessment of their impact on DNA methylation in a controlled, isogenic, and early developmental context. To further contextualize *DNMT3A*-dependent changes, we also analyzed the DNA methylation profile of hESCs engineered to harbor *NSD1* LoF mutations associated with Sotos syndrome. We focused on Sotos syndrome among overgrowth-intellectual disability syndromes due to NSD1’s role in depositing H3K36me2, a histone modification recognized by the DNMT3A PWWP domain, and prior evidence of methylation similarities between Sotos syndrome and TBRS [24], [26]. Together, these models provide a unique opportunity to dissect how DNMT3A activity is guided by histone modifications and how disruption of this interaction, through disease-associated mutations, contributes to abnormal growth phenotypes via altered DNA methylation landscapes.

## Results

### Generation of hESC models harboring *DNMT3A* and *NSD1* mutations associated with human growth syndromes

To isolate mutation-intrinsic effects from inter-individual genetic and epigenetic variability, we introduced growth syndrome-associated mutations in *DNMT3A* and *NSD1* into an isogenic H1 (WA01) hESC line (**Figure 1A**). To model TBRS, we introduced heterozygous *DNMT3A* frameshift mutations that produce premature stop codons, or the heterozygous R882H mutation, which functions in a dominant-negative manner [27], [28]. We refer to these collectively as DNMT3A LoF mutants. To model HESJAS, we introduced previously reported heterozygous W330R or D333N missense mutations in *DNMT3A* [4]. We refer to these as DNMT3A gain-of-function (GoF) mutants. For Sotos syndrome, we introduced heterozygous *NSD1* frameshift mutations that produce premature stop codons, referred to as NSD1 LoF mutants (**Table 1**). The parental cell line as well as three clones that had undergone similar targeting and selection processes without acquiring mutations in the targeted loci were used as wild type (WT) controls throughout the study.

**Figure 1.**
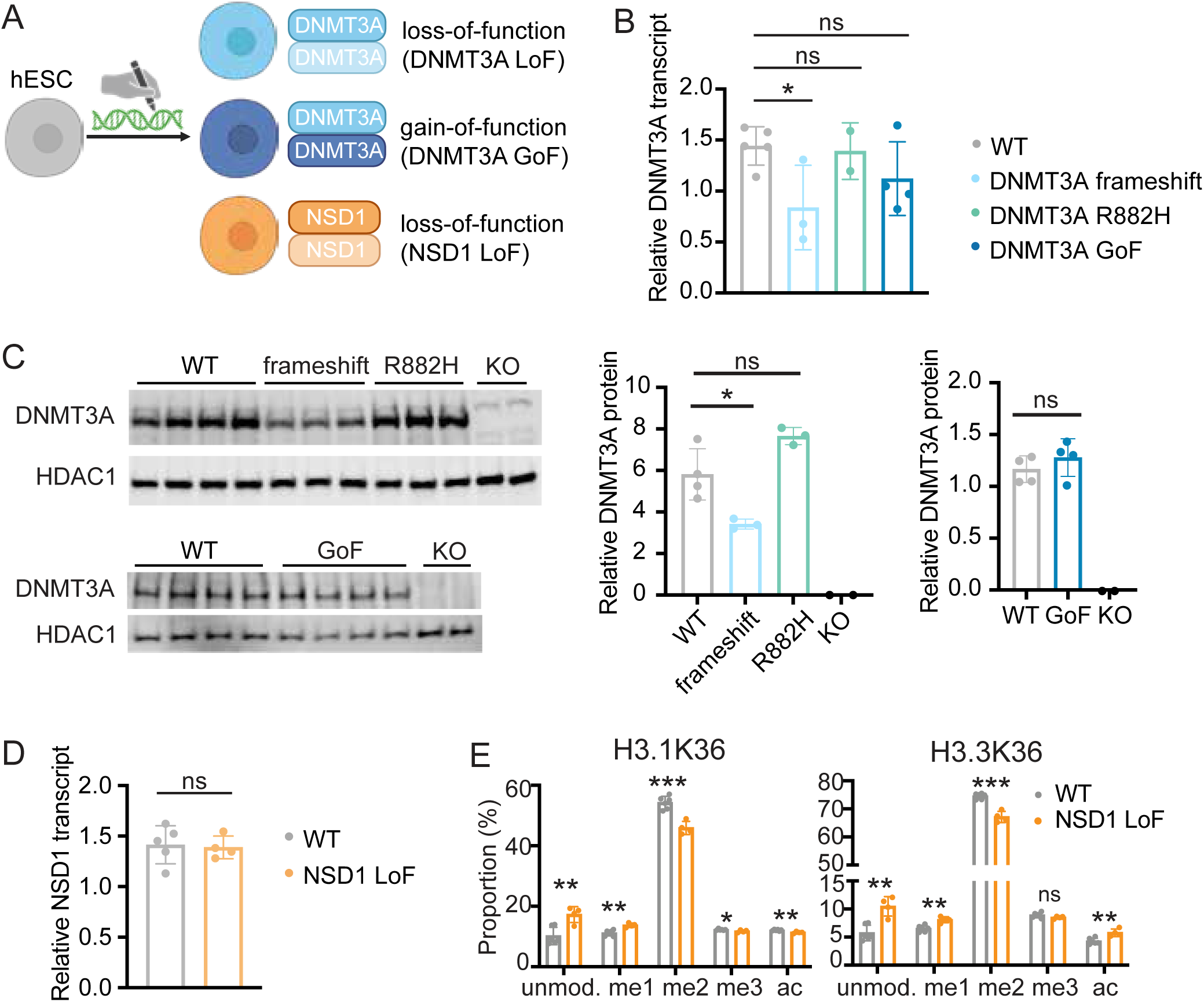
Generation and validation of hESC models for human growth syndromes. (A) Schematic of generating human embryonic stem cell (hESC) models that harbor heterozygous growth syndrome-associated mutations in *DNMT3A* and *NSD1* by CRISPR-Cas9 genome engineering. Schematic was created using BioRender. See Table 1 for detailed genotypes of the mutant clones. (B) Relative expression of DNMT3A transcripts normalized to RNA18S transcript levels. Each dot represents an independent clone. (C) Western blot analysis of DNMT3A protein expression. DNMT3A knockout (KO) clones were included as controls. HDAC1 was used as a loading control. Each lane represents an independent clone. DNMT3A genotypes are as follows: WT, WT/WT; frameshift, WT/frameshift; R882H, WT/R882H; KO, frameshift/frameshift; GoF, WT/W330R or WT/D333N. The plots on the right show the relative intensity of DNMT3A bands, normalized to HDAC1. (D) Relative expression of NSD1 transcripts normalized to RNA18S transcript levels. (E) Mass spectrometry of histones H3.1 and H3.3 K36 modifications in WT and NSD1 LoF hESCs. Each dot represents an independent clone. unmod., unmodified; me1, monomethylated; me2, dimethylated; me3, trimethylated; ac, acetylated. For panels B, C, D, and E, Statistical significance was determined by Student’s t-test. *, p<0.05; **, p<0.01; ***, p<0.001; ns, not significant.

**Table 1.** Genotype of hESC mutant clones.

| Category | Clone number | Gene | cDNA | protein |
| --- | --- | --- | --- | --- |
| DNMT3A frameshift | 1 | DNMT3A | WT/c.791_792insA | WT/p.V265Rfs*16 |
|  | 2 | DNMT3A | WT/c.791_792insTGAGC | WT/p.V265Efs*53 |
|  | 3 | DNMT3A | WT/c.989_1007del | WT/p.W330Sfs*9 |
| DNMT3A R882H | 1 | DNMT3A | WT/c.2645G>A | WT/p.R882H |
|  | 2 | DNMT3A | WT/c.2645G>A | WT/p.R882H |
| DNMT3A GoF | 1 | DNMT3A | WT/c.988T>C | WT/p.W330R |
|  | 2 | DNMT3A | WT/c.988T>C | WT/p.W330R |
|  | 3 | DNMT3A | WT/c.997G>A | WT/p.D333N |
|  | 4 | DNMT3A | WT/c.997G>A | WT/p.D333N |
| NSD1 LoF | 1 | NSD1 | WT/c.1092delC | WT/p.Y365Tfs*54 |
|  | 2 | NSD1 | WT/c.1091_1097delACTACGT | WT/p.Y364Wfs*53 |
|  | 3 | NSD1 | WT/c.1093_1094insT | WT/p.Y365Lfs*10 |
|  | 4 | NSD1 | WT/c.1092_1094delCTAinsTAGGGCCTAT | WT/p.Y365Rfs*12 |

Expression levels of pluripotency markers were comparable between mutant and WT clones (**Supplemental Figure 1**), indicating that the *DNMT3A* and *NSD1* mutations do not significantly impact hESC pluripotency under standard culture conditions. DNMT3A frameshift clones exhibited reduced mRNA and protein levels of DNMT3A compared to WT controls, presumably due to nonsense-mediate mRNA decay triggered by premature stop codons (**Figure 1B and 1C**). In contrast, DNMT3A R882H and GoF mutants maintained mRNA and protein levels of DNMT3A comparable to those in WT clones, consistent with previous reports indicating functional impairment without loss of expression [4], [28], [29] (**Figure1B and 1C**). Although NSD1 LoF mutations are predicted to introduce premature stop codons, *NSD1* mRNA levels in NSD1 LoF clones were not significantly reduced relative to those in WT controls (**Figure 1D**). To validate NSD1 LoF mutations functionally, we analyzed relative abundance of modified H3K36 via mass spectrometry. NSD1 LoF mutants displayed a significant reduction in dimethylated H3K36, with a corresponding increase in unmodified and monomethylated forms at both replication-coupled H3.1 and replication-independent H3.3 histone H3 variants [30], [31] (**Figure 1E).** These results are consistent with the reduction in H3K36me2 observed in *Nsd1*-depleted mouse ESC [6] and confirm that NSD1 LoF hESC clones are functionally impaired, despite retaining normal *NSD1* transcript levels.

### Growth syndrome associated *DNMT3A* and *NSD1* mutations cause DNA methylation defects in hESCs

We assessed DNA methylation of passage-matched mutant and WT hESCs using the Infinium MethylationEPIC BeadChip array (EPIC). After normalization and filtering, 818,992 CpG probes remained for analysis (see Methods for detail). Multidimensional scaling analysis of DNA methylation profiles separated clones according to mutation type, supporting the quality and consistency of our hESC models (**Figure 2A**). Notably, DNMT3A frameshift and R882H mutants clustered closely and showed substantial overlap in differentially methylated CpG positions (DMPs) (**Supplemental Figure 2A**). Furthermore, pairwise comparison of DMPs between frameshift and R882H clones revealed a near 1:1 correlation (**Supplemental Figure 2B**), indicating highly similar DNA methylation defects. Given their shared DNA methylation phenotype and the common clinical presentation of TBRS, we grouped them together under the DNMT3A LoF category for downstream analyses.

**Figure 2.**
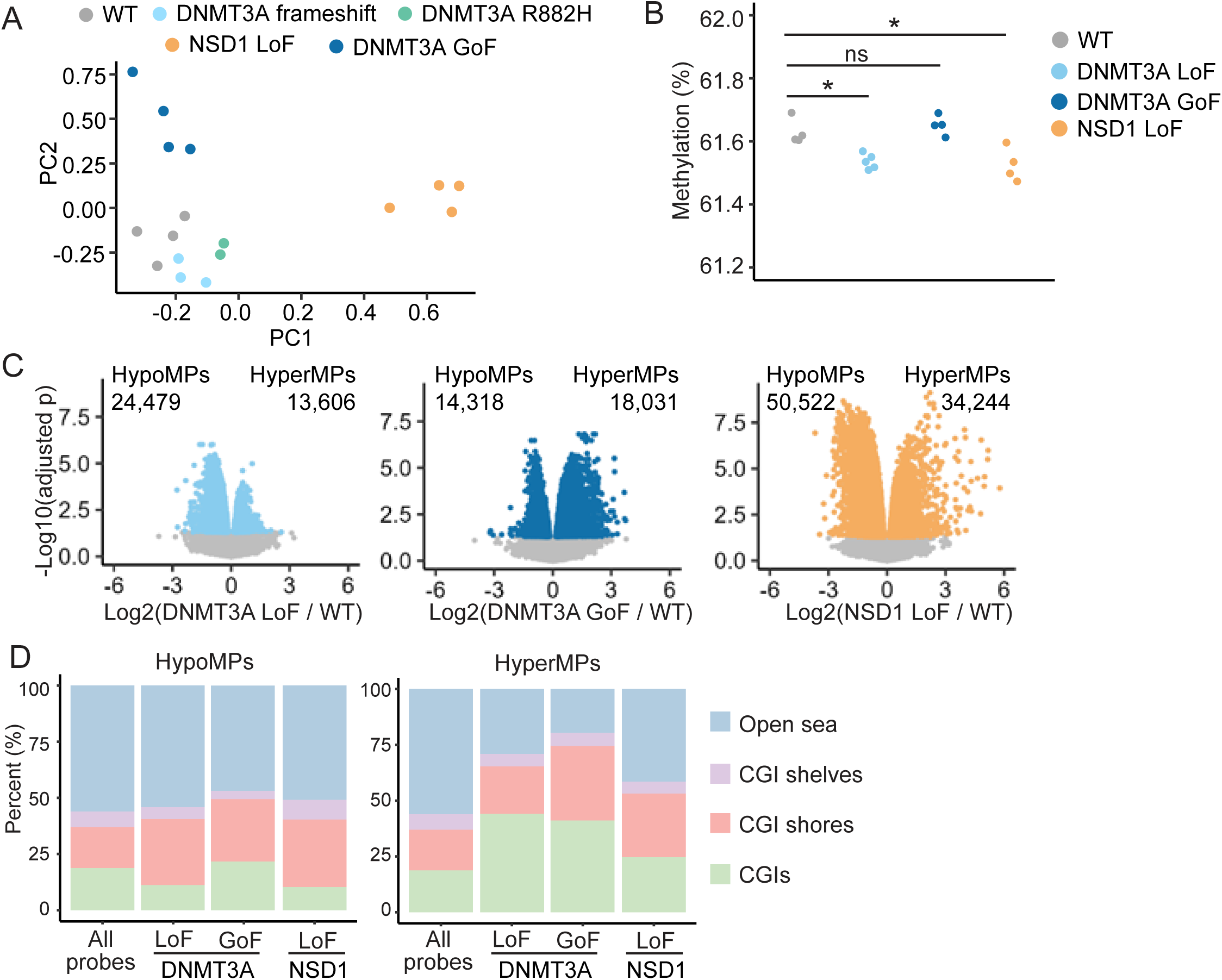
Mutant hESCs display DNA methylation defects. (A) Multi-dimensional scaling of principle components (PCs) of top 10,000 differentially methylated positions. (B) Mean methylation across all analyzed EPIC probes (n = 818,992). Each dot represents an independent clone. Statistical significance was determined by Student’s t-test. *, p<0.05; ns, not significant. (C) Volcano plots of the log2 fold change of the methylation values of mutant to WT plotted against the negative log10 of the adjusted p value. Each dot represents a probed CpG site. Colored dots, other than gray, represent significantly hypomethylated positions (HypoMPs) or hypermethylated positions (HyperMPs) with adjusted p<0.05. The number of significantly methylated positionss are indicated. (D) Distribution of HypoMPs and HyperMPs relative to CpG island (CGI) regions. CGI annotations, including CGI shores, CGI shelves, and open sea follow the definitions in the annotatr package [56], CGI shores are defined as the 2kb regions flanking the CGI boundaries CGI boundaries. CGI shelves are defined as the 2kb regions immediately upstream and downstream of the CGI shores, on the side opposite the CGIs.

The average DNA methylation across all analyzed CpG positions showed a small but statistically significant reduction in DNMT3A LoF and NSD1 LoF clones (**Figure 2B**).

Consistently, the DMP analysis revealed a greater number of hypomethylated positions (HypoMPs) compared to hypermethylated positions (HyperMPs) in these clones (**Figure 2C**). In contrast, DNMT3A GoF clones exhibited more HyperMPs than HypoMPs, despite their overall average DNA methylation not significantly differing from WT controls (**Figure 2B and 2C**). To further characterize these differences, we examined the genomic distribution of DMPs relative to CpG islands (CGIs) (**Figure 2D**). HypoMPs across all mutant categories showed enrichment at CpG island (CGI) shores, while HyperMPs were enriched with CGIs. These significant methylation changes at CGIs and their vicinity suggested that we are capturing biologically meaningful DNA methylation changes.

### DNA methylation defects in mutant hESCs mirror those observed in fully differentiated patient cells

We next examined whether the DNA methylation defects in mutant hESC lines reflect those present in patient samples. DNA methylation data from blood of individuals with DNMT3A- or NSD1-associated growth syndromes, generated using the Infinium HumanMethylation450K or EPIC arrays, were obtained from published studies [32], [23], [4] (**Supplemental Table 1).** To compare these datasets, we annotated each DMP using a chromatin state model which classifies genomic regions based on shared histone modification patterns, enabling genome-wide analysis with reduced dimensionality [33], [34]. To be able to compare our hESC data to patient methylation data from blood samples, we used a tissue-independent full-stack chromatin state annotation. This framework defines a set of broad chromatin states and further subdivides them into cell type–specific substates, resulting in a total of 100 distinct chromatin states [34].

The predominant DMP category of each hESC mutant (HypoMPs for DNMT3A LoF and NSD1 LoF, and HyperMPs for DNMT3A GoF) displayed chromatin state enrichment patterns strikingly similar to those observed in the matching DMP category of corresponding patient blood samples (**Figure 3**). HypoMPs from DNMT3A LoF hESCs and TBRS patient blood were significantly enriched in enhancer (EnhA), bivalent promoter (BivProm), and promoter-flanking (PromF) chromatin states (**Figure 3A and 3B**), while repressed Polycomb (ReprPC) and bivalent promoter states were enriched in HyperMPs from DNMT3A GoF hESCs and HESJAS patient blood (**Figure 3C and 3D**). HypoMPs of NSD1 LoF hESCs and Sotos syndrome patient blood showed similar enrichment patterns compared to those in DNMT3A LoF hESCs and TBRS blood, with an additional enrichment in repressed Polycomb states (**Figure 3E and 3F**). Together, these results demonstrate that DNA methylation defects arise even in undifferentiated pluripotent stem cells harboring disease-associated mutations. Furthermore, the hESC models recapitulate DNA methylation defects observed in patient cells, indicating that the chromatin states affected by these mutations are conserved between pluripotent and fully differentiated cells.

**Figure 3.**
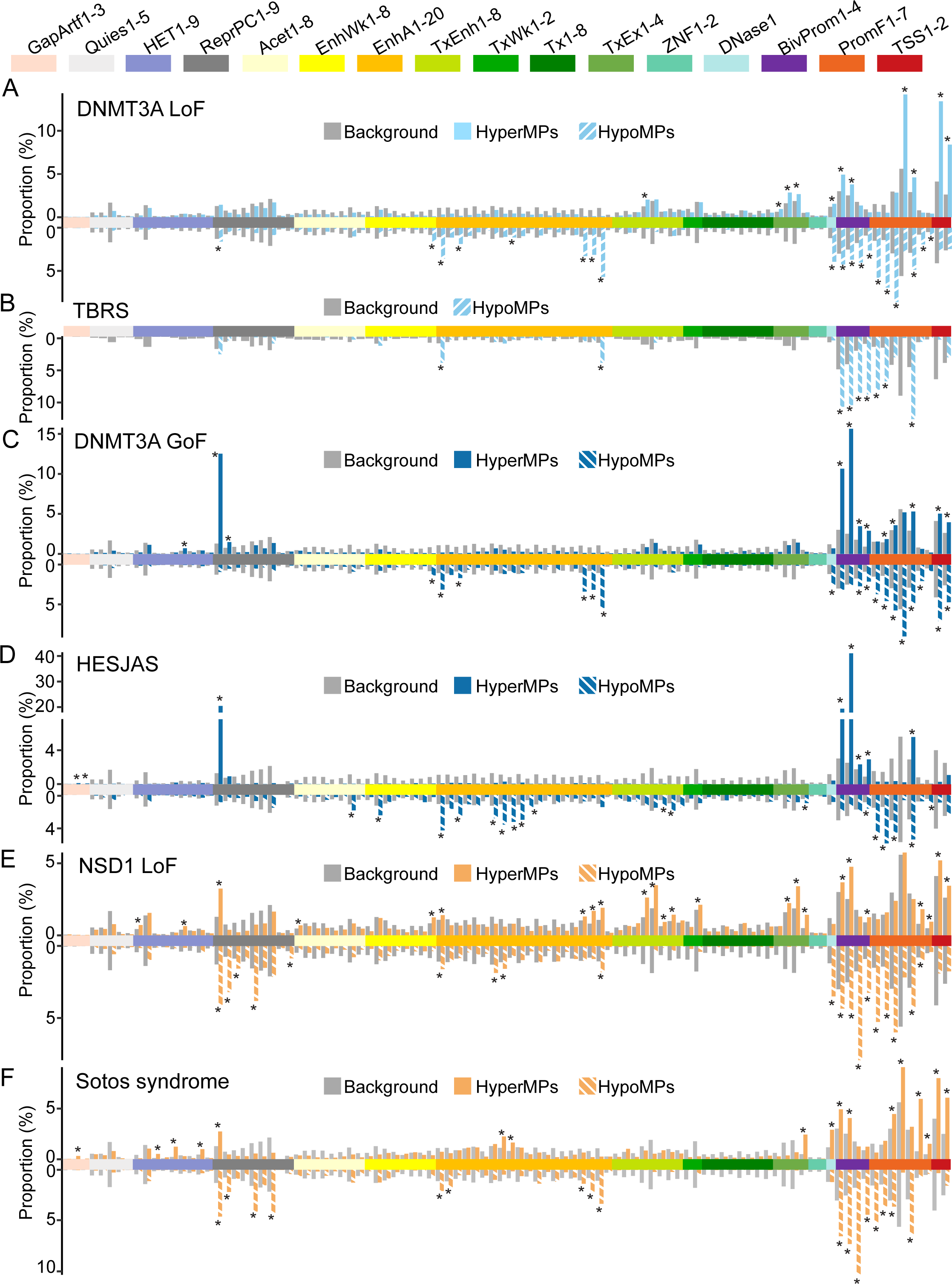
Chromatin state analysis of differentially methylated positions in hESC mutants and patient blood. Proportions of in each full-stack chromatin state for (A) DNMT3A LoF hESCs, (B) TBRS patient blood (HypoMPs n=832), (C) DNMT3A GoF hESCs, (D) a HESJAS patient peripheral blood leukocyte sample (HypoMPs n=2,796; HyperMPs n=9,576), (E) NSD1 LoF hESCs, and (F) Sotos syndrome patient blood (HypoMPs n=24,148; HyperMPs n=4,168) Background represents proportion of all probes in each state. HyperMPs of TBRS patients were not analyzed due to a small size (n=38). Statistical significance was determined by Fisher test. *, p<0.001.

Legend of cell-type independent chromatin states [34]: GapArtf, assembly gaps and alignment artifacts; Quies, quiescent (low histone emissions, except possibly weak H3K9me3); HET, heterochromatin (H3K9me3); ReprPC, repressed polycomb (H3K27me3); Acet, various acetylations (with weaker H3K4me1/2/3, H3K9ac, and H3K27ac emissions); EnhWk, weak enhancers; EnhA, active enhancers (enhancers marked by H3K4me1, DNase, H2A.Z, and/or H3K27ac); TxEnh, transcribed candidate enhancers; TxWk, weak transcription; Tx, strong transcription; TxEx, transcription and exons (transcription marked by H3K36me3, H3K79me1, H3K79me2, or H4K20me1); ZNF, zinc finger (H3K36me3 and H3K9me3); DNase1, only DNase 1 hypersensitivity; BivProm, bivalent promoter (H3K27me3 and promoter marks); PromF, promoter flanking (H3K4me1 and promoter marks); TSS, transcriptional start sites (promoter marks of H3K4me2/3 and H3K9ac).

### H2AK119ub-marked active enhancers are hypomethylated in both DNMT3A LoF and GoF mutants, revealing a role of the PWWP-H3K36me2/3 interaction in maintaining DNA methylation at these regions

Although fewer than HyperMPs, DNMT3A GoF hESC mutants as well as HESJAS patient blood exhibit a substantial number of HypoMPs (**Figure 2C, 3C and 3D**). We were intrigued by the highly similar enrichment pattern of chromatin states of DNMT3A GoF HypoMPs and that of DNMT3A LoF HypoMPs (**Figure 3A and 3C)**. While the hypomethylation phenotype of DNMT3A LoF and TBRS patients has been extensively studied [23], [24], [25], [35], previous studies of DNMT3A GoF mutations have focused mainly on hypermethylated regions with limited characterization of hypomethylated ones [4], [36].

DNMT3A GoF mutations in HESJAS disrupt the PWWP domain’s ability to bind H3K36me2/3 without affecting overall protein levels [4], [7], [37] (**Figure 1C**), whereas DNMT3A LoF mutations broadly diminish the availability of catalytically active DNMT3A at its methylation targets either via reduced protein levels (frameshift) or impaired catalytic activity (R882H). Thus, we hypothesized that HypoMPs shared between DNMT3A GoF and DNMT3A LoF mutants represent genomic regions whose methylation relies on the DNMT3A PWWP-H3K36me2/3 interaction. To identify such regions, we intersected HypoMPs from DNMT3A LoF mutants with those from DNMT3A GoF mutants and analyzed chromatin states enriched at these shared sites (shared HypoMPs) (**Figure 4A)**.

**Figure 4.**
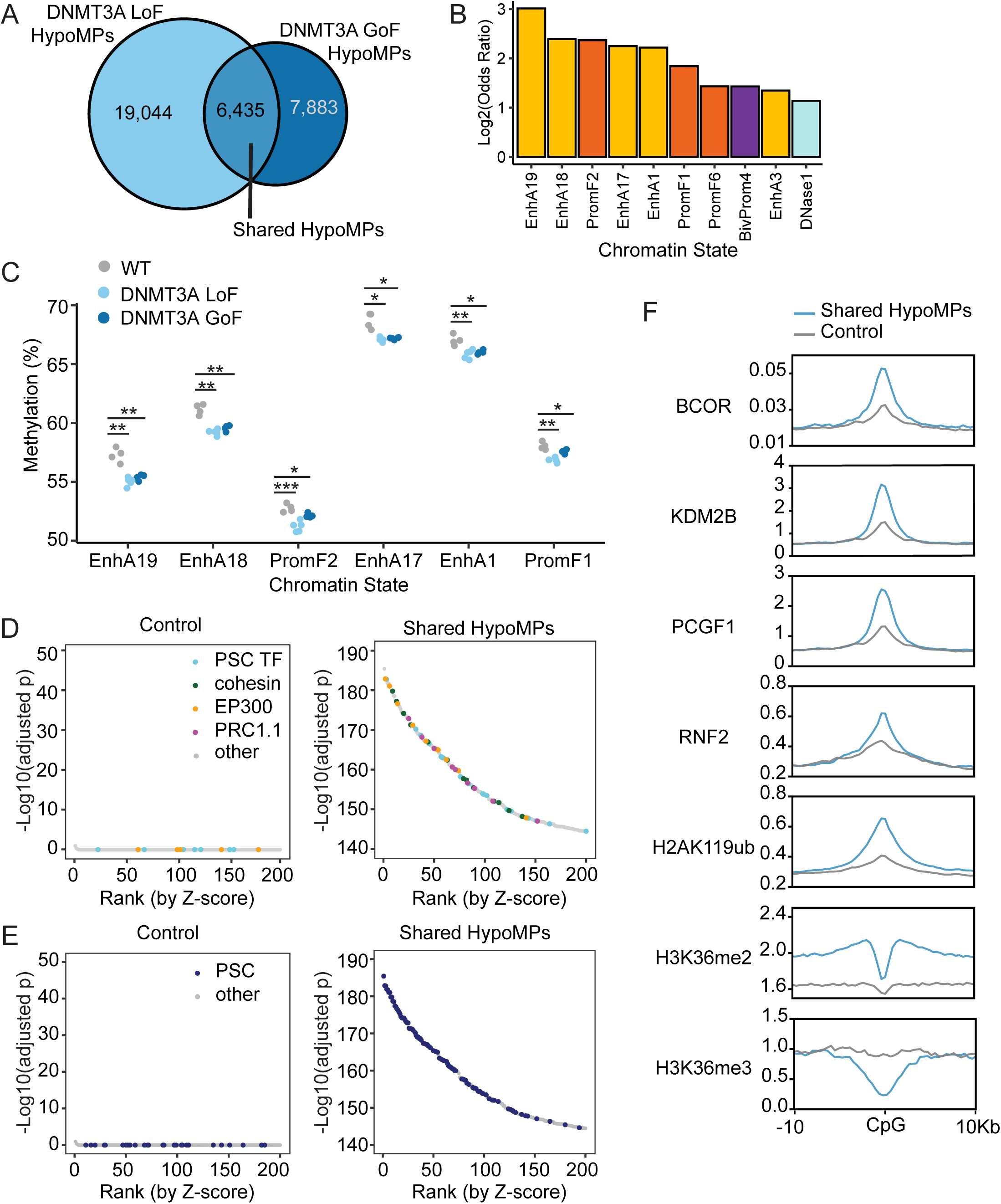
Shared hypomethylation of H2AK119ub-marked active enhancers in DNMT3A GoF and LoF mutants. (A) Number of overlapping HypoMPs between DNMT3A LoF and GoF mutants. P<2.2*10^-16, Fisher’s exact test of unique DNMT3A GoF in all probes compared to DNMT3A GoF shared with DNMT3A LoF. (B) Log2 odds ratios showing enrichment of shared HypoMPs from DNMT3A mutants across full-stack chromatin states. Top 10 enriched states are shown (all p<1*10-18). (C) Mean DNA methylation values at CpG positions within selected enhancer chromatin states. Each dot represents an independent clone. Statistical significance was determined by Student’s t-test. *, p<0.05; **, p<0.01; ***,p<0.001; ns, not significant. (D-E) RELI analysis of shared HypoMPs intersected with >10,000 public chromatin datasets. Shown are the top 200 datasets ranked by Z-score. Most enriched chromatin factors (D) and cell types (E) are highlighted. PSC TF: POU5F1, NANOG, SOX2, cohesion: RAD21, NIPBL, PRC1.1: BCOR, KDM2B, PCGF1, RYBP, RNF2. Controls represent randomly sampled EPIC probe regions. Each dot represents a dataset profiling a chromatin factor in a human cell type. Supplemental Table 2 contains RELI results of all examined datasets. PSC, pluripotent stem cells; TF, transcription factors. (F) CUT&RUN (H2AK119ub) or ChIP-seq (all others) signal in WT hESCs 10 kilobases upstream and downstream from the center of shared HypoMPs or matched control regions. BCOR, KDM2B, PCGF1, H3K36me2 occupancy data are from GEO accession number GSE104690, RNF2 data is from GSE105028, H3K36me3 is from ENCODE (ENCSR476KTK), and H2AK119ub is from GSE301386 (this study).

The shared HypoMPs were significantly enriched in several enhancer and promoter-flanking chromatin states (**Figure 4B)**. Consistently, pronounced reduction in DNA methylation was observed in enhancer states EnhA 1,17,18, and 19 in both DNMT3A LoF and DNMT3A GoF mutants (**Figure 4C**). While EnhA1 corresponds to enhancers broadly active across cell types, EnhA 17-19 are specific to pluripotent stem cells including hESCs [34]. Interestingly, HypoMPs in the HESJAS patient’s blood were more strongly enriched in blood lineage-specific enhancer states (EnhA7-11) than in pluripotency-specific ones **(Figure 3D**). Together, these findings suggest that the DNMT3A PWWP domain is required for maintaining DNA methylation at enhancers active in the corresponding cell type.

To further characterize the shared HypoMPs, we performed a Regulatory Element Locus Intersection (RELI) analysis by intersecting the shared HypoMPs with over 10,000 publicly available occupancy maps of chromatin-associated proteins across diverse human cell types.

RELI revealed highly significant overlap between shared HypoMPs and chromatin regions occupied by pluripotency-associated transcription factors (e.g., POU5F1, NANOG, SOX2), cohesion complex components and loader (e.g. RAD21, NIPBL), and transcriptional coactivators, particularly EP300 (**Figure 4D, Supplemental Table 2**). Notably, the vast majority of the most significantly overlapping datasets were generated in pluripotent stem cells (**Figure 4E, Supplemental Table 2**). Together, these results independently validate that shared HypoMPs are enriched in pluripotency-specific active enhancers.

RELI also uncovered significant overlap between shared HypoMPs and genomic regions occupied by PRC1.1 components, including BCOR, KDM2B, PCGF1, and RNF2 (**Figure 4D and 4F)**. PRC1.1 is a noncanonical variant of PRC1 that deposits H2AK119ub1 [38], and accordingly, we observed H2AK119ub1 enrichment at shared HypoMPs compared to control CpG sites (**Figure 4F**). This was unexpected, as previous studies have shown that DNMT3A GoF mutant proteins, which lack the ability to bind H3K36me2/3, are redirected to H2AK119ub1-marked regions via the N-terminal ubiquitin-dependent recruitment motif, resulting in hypermethylation [8], [37]. Our data indicate that a subset of H2AK119ub1-marked regions, particularly enhancers embedded in environments enriched for H3K36me2 but not H3K36me3, are hypomethylated in DNMT3A mutants (**Figure 4F**). This suggests that recognition of H3K36me2 by the DNMT3A PWWP domain is required for proper DNA methylation at these sites, and supports H3K36me2 as the primary histone mark guiding DNMT3A recruitment in this context.

### Mutations associated with overgrowth versus growth restriction cause contrasting DNA methylation changes at bivalent chromatin

Our data suggest that while hypomethylation of enhancers is a prominent feature in DNMT3A LoF mutants and TBRS, it is unlikely to underlie the overgrowth phenotype, as the same pattern is also present in DNMT3A GoF mutants and HESJAS patient with growth restriction. To investigate alternative DNA methylation signatures potentially related to growth regulation, we searched for genomic regions with shared methylation changes in the two overgrowth-associated conditions—DNMT3A LoF and NSD1 LoF—but with opposing methylation changes in DNMT3A GoF mutants.

Direct CpG-level comparison identified only 156 CpG sites that were significantly hypomethylated in both DNMT3A LoF and NSD1 LoF mutants and hypermethylated in DNMT3A GoF mutants, with no CpGs exhibiting the reverse pattern. This limited overlap suggested that growth-related divergent methylation changes might occur at the chromatin state level rather than individual CpG sites. We noted that bivalent promoter states (BivProm1-4) were markedly enriched among HyperMPs in DNMT3A GoF mutants, while enriched among HypoMPs in DNMT3A LoF and NSD1 LoF mutants (**Figure 3A, 3C, 3E**). Importantly, these chromatin-state-specific patterns were mirrored in patient blood samples: bivalent promoter states were hypermethylated in growth-restricted HESJAS patients and hypomethylated in overgrowth syndromes TBRS and Sotos (**Figure 3B, 3D, 3F**).

Bivalent promoters, defined by co-occupancy of H3K4me3 and H3K27me3, are typically associated with developmental genes that are poised for activation, but show low transcriptional activity in pluripotent stem cells [39]. To characterize these regions further, we clustered regions marked by H3K4me3 in WT H1 based on their H3K27me3 signal intensity (**Figure 5A**). As expected, H3K27me3 levels inversely correlated with H3K4me3 levels. We also observed a positive correlation between H3K27me3 signal strength and the degree of DNA hypermethylation in DNMT3A GoF mutants (**Figure 5B**). Accordingly, clusters 1–3 showed significant hypermethylation in DNMT3A GoF mutants, while cluster 4, with the weakest H3K27me3 signal, did not. Notably, only cluster 3 regions, characterized by relatively weak levels of both H3K4me3 and H3K27me3, were significantly hypomethylated in both DNMT3A LoF and NSD1 LoF mutants (**Figure 5B**). These results suggest that altered DNA methylation at bivalent promoters, with balanced low levels of H3K4me3 and H3K27me3, may contribute to growth dysregulation in DNMT3A- and NSD1-related overgrowth and growth restriction syndromes.

**Figure 5.**
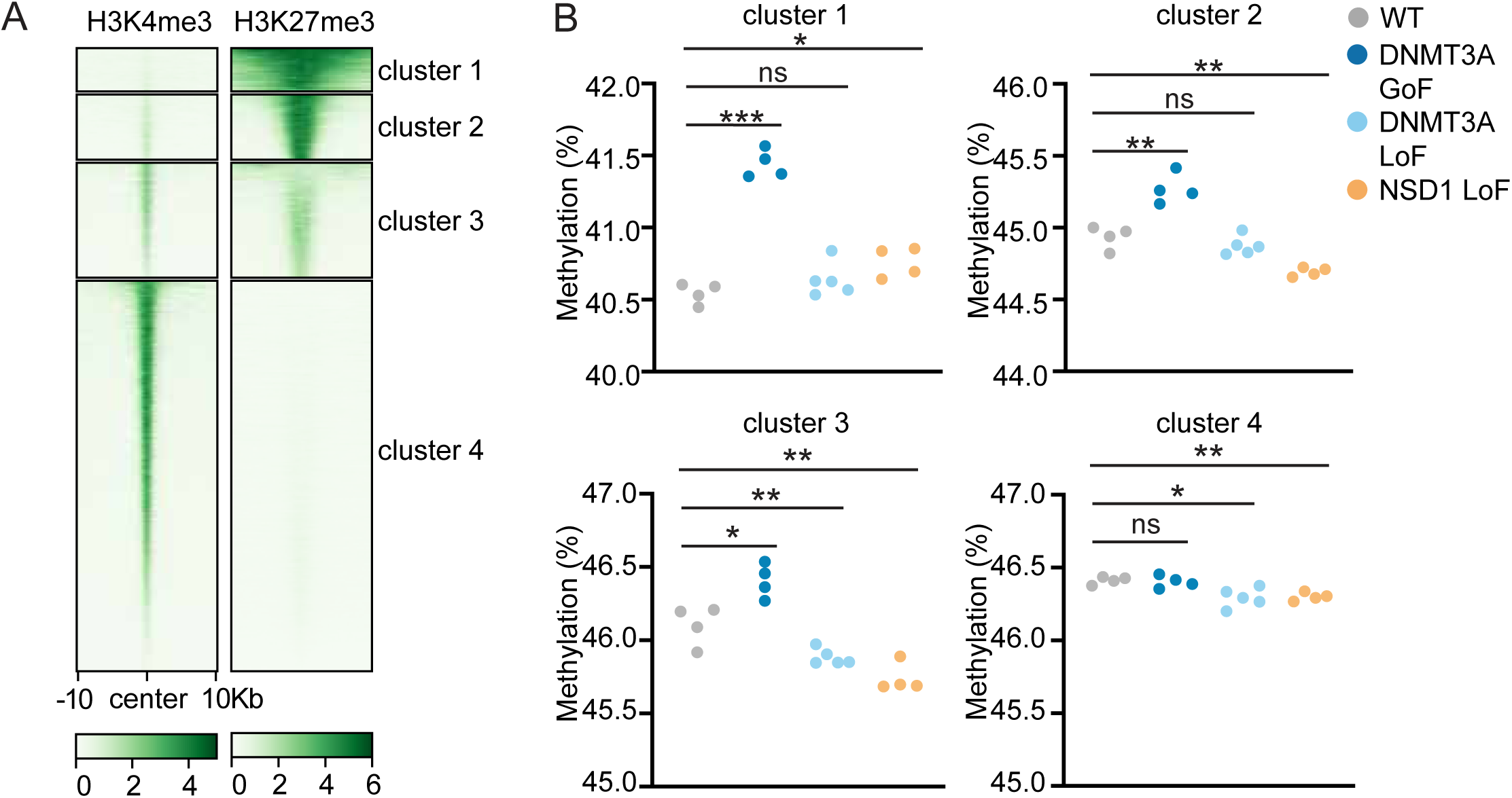
Contrasting DNA methylation changes at bivalent chromatin in overgrowth versus growth restriction hESC models. (A) Heatmap of H3K4me3 and H3K27me3 signals at H3K4me3-marked genomic regions, clustered based on H3K27me3 signal intensity in WT H1 hESCs. (B) Mean DNA methylation values at CpG positions within the indicated clusters from panel (A). Each dot represents an independent clone. Statistical significance was determined by Student’s t-test. *, p<0.05; **, p<0.01; ***, p<0.001; ns, not significant.

## Discussion

In this study, we compared shared and distinct molecular defects in hESCs engineered with *DNMT3A* and *NSD1* mutations that underlie overgrowth and growth restriction syndromes. Our models recapitulate DNA methylation changes observed in patient blood samples, as confirmed by reanalysis of publicly available datasets, which is consistent with published interpretations [4], [23], [24], [25], [32], [35], [36]. These similarities underscore the utility of hESCs as a model system for investigating the molecular pathology of these disorders.

Interestingly, the reduction in H3K36me2 in NSD1 LoF hESCs was less than 50%, suggesting compensation by related histone methyltransferases such as NSD2 and NSD3, as recently shown [40]. Likewise, DNMT3A haploinsufficiency does not halve DNA methylation, likely due to both overlapping activity from DNMT1 and DNMT3B and the possibility that DNMT3A protein levels from a single functional allele are sufficient to maintain DNA methylation at most loci. Despite these compensatory mechanisms and near-normal global modification levels, disease still develops, indicating that specific genomic regions are critically dependent on the full dosage of *NSD1* or *DNMT3A*.

By introducing mutations into an isogenic background, we were able to directly compare mutation-specific effects without the confounding variables present in patient samples. This approach revealed an unexpected finding: shared hypomethylation at H2AK119ub-marked active enhancers in both DNMT3A GoF and LoF mutants. This suggests a cooperative role of H2AK119ub and H3K36me2/3 in recruiting DNMT3A to active enhancers, and not just a competitive one [37]. Although this shared enhancer hypomethylation phenotype is unlikely to explain the divergent growth outcomes in TBRS and HESJAS, its functional relevance warrants further investigation, as it may contribute to other syndromic features associated with these disorders.

The contrasting DNA methylation defects observed in overgrowth versus growth restriction mutants implicate bivalent chromatin as a potential regulatory hub in organismal growth. However, the relationship is unlikely to follow a simple inverse correlation between DNA methylation and gene expression (i.e. hypomethylation leading to gene upregulation and vice versa). For instance, in blood samples from Sotos patients with overgrowth syndrome, genes associated with bivalent chromatin were downregulated, contrary to the upregulation typically expected from hypomethylation [32]. This underscores the need for careful analyses of gene expression across different stages of differentiation to better understand how DNA methylation defects influence transcriptional programs of bivalent genes and ultimately contribute to growth outcome in DNMT3A- and NSD1-related disorders.

Regulation of bivalent chromatin, particularly through modulation of PRC2 activity and H3K27me3 levels, appears to play a broader role in other overgrowth and growth restriction syndromes. For example, partial LoF mutations in EZH2, the catalytic subunit of PRC2, are associated with Weaver syndrome, the second most prevalent overgrowth-intellectual disability syndrome [11], [19]. These mutations have been shown to impair histone methyltransferase activity and reduce methylated H3K27 [41], [42], [43]. Conversely, a missense EZH2 mutation p.A738T, which enhances its catalytic activity [44], [45], has been reported in a patient with growth restriction [44], suggesting that the levels of H3K27me3 modulate growth outcomes in a dosage-dependent manner. Given the documented crosstalk among DNMT3A, NSD1, and PRC2 [46], it would be informative to compare gene expression changes across these models to identify shared or divergent regulatory consequences.

Our hESC models complement patient data while allowing further molecular characterization and manipulation, which future studies will explore in differentiated cell types and map alongside gene expression changes. Finally, our findings reinforce the role of PRC-regulated regions as key methylation targets in growth syndromes and underscore the importance of understanding PRC-DNMT interactions in both normal development and disease.

## Methods

### Generation of hESCs mutant lines

The hESC line H1 (WA01) was obtained from WiCell and maintained under feeder-free condition in StemFlex media (Thermo Fisher Scientific) on Cultrex (R&D Systems)-coated plates. Cells were passaged using Accutase (STEMCELL Technologies) and replated in StemFlex medium supplemented with 1 μM thiazovivin (Selleck Chemicals) to enhance viability. To introduce mutations associated with the growth syndromes, synthetic guide RNAs (**Supplemental Table 3**) (Integrated DNA Technologies) were complexed with recombinant Streptococcus pyogenes Cas9 nuclease (Integrated DNA Technologies) to form ribonucleoprotein (RNP) complexes. These RNPs were delivered into H1 cells by electroporation using either the Neon Transfection System (Thermo Fisher Scientific) or the 4D-Nucleofector (Lonza). For DNMT3A R882H and DNMT3A GoF mutations, a synthetic single-stranded oligonucleotide (ssODN) donor template (**Supplemental Table 3**) (Integrated DNA Technologies) was co-transfected to facilitate homology-directed repair. Following a 48-hour recovery period, transfected H1 cells were replated onto irradiated mouse embryonic fibroblasts (iMEFs) (Thermo Fisher Scientific) to allow for the formation of single-cell-derived colonies. After 10–14 days, individual colonies were picked and genotyped by PCR amplification of genomic regions flanking the target sites, followed by Sanger sequencing to identify clones carrying the desired mutations. At least three clones per mutation category (WT, DNMT3A LoF, DNMT3A GoF, and NSD1 LoF), each derived from separate wells, were subjected to an additional round of single-cell cloning on iMEFs to ensure clonality and eliminate mixed populations prior to downstream applications.

### Real-time quantitative PCR

Total RNA was extracted using Quick-RNA Miniprep Kit (Zymo Research) according to the manufacturer’s protocol. A total of 500 ng of RNA was used for reverse transcription using High-Capacity cDNA Reverse Transcription Kit (Thermo Fisher Scientific). RT-qPCR reactions were prepared using 6.25 ng of total cDNA in a total volume of 10 μL with PowerUp SYBR Green Master Mix (Thermo Fisher Scientific), following the manufacturer’s instructions. Amplification was performed on CFX384 Touch Real-Time PCR Detection System (Bio-Rad Laboratories). All reactions were performed in triplicate and melt curve analysis was used to confirm specificity.

Gene-specific primers targeting human DNMT3A (5’-CCTCTTCGTTGGAGGAATGTGC-3’, 5’-GTTTCCGCACATGAGCACCTCA-3’), NSD1 (5’ -CAAGGAAGCGAAAACGACAGAGG-3’, 5’-CCGTCCTGTGAGGCATTAGTTC-3’), and 18S ribosomal RNA (RNA18S) (5’-ACCCGTTGAACCCCATTCGTGA-3’, 5’-GCCTCACTAAACCATCCAATCGG-3’) as an endogenous control were used for this assay. Relative gene expression levels were calculated using the ΔCt method, with normalization to RNA18S.

### Western blot

Cells were washed with calcium- and magnesium-free PBS and lysed in 4× Laemmli sample buffer (Bio-Rad Laboratories) containing 2-mercaptoethanol. Lysates were boiled at 95 °C for 5 minutes. Protein concentrations were measured using Pierce’s 660 nm Protein Assay Reagent (Thermo Fisher Scientific) according to the manufacturer’s instructions. Equal amounts of protein (10 μg) were loaded onto 4–20% Mini-PROTEAN TGX Stain-Free Gels (Bio-Rad Laboratories) and electrophoresed. Proteins were transferred onto PVDF membranes using semi-dry transfer in buffer containing 20% ethanol. Membranes were blocked with EveryBlot Blocking Buffer (Bio-Rad Laboratories) for 10 minutes at room temperature. Membranes were incubated overnight at 4 °C with a primary antibody against DNMT3A (C-12, Santa Cruz Biotechnology, sc-365769), followed by HRP-conjugated anti-mouse IgG secondary antibody (Cell Signaling Technology, #7076) for 1 hour at room temperature. HDAC1 (10E2, Cell Signaling Technology, #59581) was used as a loading control. Between steps, membranes were washed 3 × 5 minutes with TBST. Immunoreactive bands were visualized using ECL (Thermo Fisher Scientific) and imaged with ChemiDoc imaging system (Bio-Rad Laboratories).

### Flow cytometry

hESCs were dissociated into single-cell suspensions using 0.05% Trypsin-EDTA (Thermo Fisher Scientific) and stained with LIVE/DEAD™ Fixable Dead Cell Stain Sampler Kit (Thermo Fisher Scientific) according to the manufacturer’s instructions. Cells were washed with calcium- and magnesium-free PBS supplemented with 5 mM EDTA and 2% FBS and then incubated with APC-conjugated anti-CD326 (EpCAM) monoclonal antibody (clone 323/A3, Thermo Fisher Scientific, MA5-38715) or mouse IgG1 kappa isotype control antibody at a density of one million cells per ml for 30 minutes at 4 °C in the dark. After staining, cells were washed and analyzed using NovoCyte Quanteon Flow Cytometer System (Agilent Technologies). Data was acquired and analyzed with FlowJo software (BD Biosciences).

### Mass spectrometry

For sample preparation, culture media was removed, and cells were washed twice with Dulbecco’s Phosphate-Buffered Saline (DPBS). Following DPBS removal, cells were scraped and collected into 1.5 mL microcentrifuge tube, and centrifuged at 2000g for 3 minutes. The supernatant was discarded, and the resulting cell pellet was immediately snap-frozen in liquid nitrogen and stored at -80 °C until further processing. Histones were acid-extracted, derivatized via propionylation, and digested with trypsin, as previously described [47]. Each sample was resuspended in 300 µL of 0.1% FA/mH2O, and 2 µl was injected per run, with 3 technical replicates. Histone extraction, histone modification profiling, and mass spectrometry analysis were conducted at the Mass Spectrometry Technology Access Center at Washington University School of Medicine.

### DNA methylation array and analysis

Genomic DNA was extracted from hESCs using Quick-DNA Kit (Zymo Research). DNA methylation levels were assessed with Infinium MethylationEPIC BeadChip array (Illumina) according to the manufacturer’s instructions. DNA methylation data was processed in R (v4.4.0) based on Bioconductor workflow by Maksimovic and colleagues with minor alterations [48], [49]. In brief, Infinium™ idat files were processed and mapped to human genome 38 (hg38) using preprocessQuantile in minfi (v1.50.0) [50]. Probes for non-CpG sites, probes not mapped to hg38, and poor quality probes with detection p values greater than 0.01 were removed. Cross reactive probes were removed using maxprobes package (v0.0.2) [51]. For data obtained from patients (**Supplemental Table 1**), probes with common SNPs at the CpG site or at single nucleotide base extensions were removed (minfi). Sex chromosome probes were removed in patient datasets with males and females. M-values and beta-values were then extracted and M-values were adjusted with ComBat for individual BeadChip batch effects if multiple chips were used (sva v3.52.0) [52], [53]. Adjusted M-values were fit to a linear model and tested with an empirical Bayes test with a Benjamini & Hochberg (FDR) correction for multiple comparisons using limma (v3.60.6) [54]. CpG sites with an adjusted p-value <0.05 were extracted for analyses involving DMPs. Batch corrected M-values were coerced back to beta values by taking the inverse log2 and were reported as methylation percentages. Chromatin state regions were downloaded as bed files and overlapping DMPs were summarized using annotatr (v1.30.0) and genomic ranges (v1.56.2) [55], [56]. DNA methylation plots were generated in R using ggpolot2 (v3.5.1), ggpubr (v0.6.0.999), and ggbreak (v0.1.4) [57], [58], [59] or in GraphPad Prism (v10.1.0) (ref).

### RELI

We collected 10,709 publicly available functional genomic datasets from the Gene Expression Omnibus repository [60] and processed them uniformly using the ENCODE ChIP-seq pipeline [61]. Final ChIP-seq peak sets were indexed by their hg38 genomic coordinates and intersected with the coordinates of DNA methylation sites of interest. To assess the statistical significance of these overlaps, we used RELI [62], which compares observed intersections to a null model. The null model was generated by padding all analyzed CpG sites (818,992 in total) with 100 bp upstream and downstream, then merging overlapping regions to yield 531,738 non-redundant genomic intervals. The distribution of the expected intersection values from the null model resembles, and therefore was modeled as, a normal distribution, where the model parameters were estimated from a series of sampling procedures using RELI. The significance of the observed number of intersections, e.g., a Z-score and the corresponding P-value, was calculated by applying the same padding and merging strategy to DMPs and comparing their overlap with ChIP-seq peaks to the null distribution. As a control, we randomly sampled the same number of genomic intervals from the null model’s genomic intervals.

### CUT&RUN

CUT&RUN was performed using the CUTANA CUT&RUN Kit (Cat # 14-1048, EpiCypher, NC, USA) and targeted histone mark antibodies (anti-H3K4me3, anti-H3K27me3, EpiCypher, NC, USA; anti-H2AK119ub, Cell Signaling Technology, MA, USA) following the manufacturer’s protocol. In brief, 0.5 million cells were harvested and incubated with activated Concanavalin A for 10 min at room temperature. For each target histone mark, 0.5 μg of H3K4me3 (Cat # 13-0041), H3K27me3 (Cat # 13-0030), or H2AK119ub (Cat # 8240) was added to each sample and incubated overnight at 4 °C. Isotype control (Cat # 13-0042, EpiCypher) was used as a negative control. After overnight incubation, the beads were then washed twice with Cell permeabilization buffer (Wash buffer including 0.01% digitonin), and incubated with protein AG-Micrococcal Nuclease (pAG-MNase) for 1h at 4 °C. Excessive pAG-MNase was washed out, then chromatin digestion was performed by adding 2mM CaCl2. After chromatin digestion, the stop buffer (Cat # 48105, Cell signaling Technology) and 1 ng *E.coli* spike-in DNA (Cat # 18-1401, EpiCypher) were added and incubated for 10min at 37 °C.

CUT&RUN libraries were prepared with the CUT&RUN library Prep kit (Cat# 14-1001,, EpiCypher) following the manufacturer’s instructions. Libraries were quantified with the Quant-iT picogreen dsDNA assay kit (Cat# P7589, Thermo Fisher Scientific) and fragment sizes assessed using an Agilent 2100 Bioanalyzer system. Libraries were sequenced to have at least 6 million read pairs using an Illumina NovaSeq 6000. The raw sequencing reads were first trimmed with TrimGalore v0.6.7 and then aligned to the human reference genome hg38 (GRCh38) using Bowtie2 v2.5.1 (parameters --local --very-sensitive-local --no-unal --no-mixed --no-discordant --phred33 -I 10 -X 700). Sam files were sorted and indexed with samtools v1.15.1 to generate bam files, and then duplicate reads were removed using gatk markduplicates (v4.2.6.1). After removing PCR duplicates and unaligned reads, bigWig files were generated from bam file using bedtools2 v2.30 and reads were normalized by total counts per sample.

Heatmap and histone mark enrichment distribution were created using deepTools [63].

### Clustering of H3K4me3-marked regions

H3K4me3 ChIP-seq peaks in H1 hESCs were obtained from ENCODE (ENCSR443YAS). Using the deepTools [63] computeMatrix function in reference-point mode, H3K4me3 and H3K27me3 signals from our CUT&RUN data (GEO accession: GSE301386) were quantified, centered on peak center, and extended from the center ±10 kb with a bin size of 400 bp. K-means clustering (k = 4) was performed based on the H3K27me3 signal profile across these regions.

## Abbreviations

CGI: CpG island
CpG: cytosine followed by guanine
DMPs: differentially methylated CpG positions
GoF: gain-of-function
hESC: human embryonic stem cell
HESJAS: Heyn-Sproul-Jackson syndrome
HyperMPs: hypermethylated positions
HypoMPs: hypomethylated positions
LoF: loss-of-function
PRC: Polycomb Repressive Complex
PWWP: Pro-Trp-Trp-Pro
RELI: Regulatory Element Locus Intersection
TBRS: Tatton-Brown-Rahman syndrome
TF: transcription factor

## Data availability

The DNA microarray have been deposited in GEO (accession: GSE299394). Patient DNA methylation array GEO accession numbers are in **Supplemental Table 1**. BCOR, KDM2B, PCGF1, H3K36me2 occupancy data are from GEO accession number GSE104690, H3K36me3 is from ENCODE (ENCSR476KTK), RNF2 data are from GSE105028, and H2AK119ub is from GSE301386 (this study).

## Supplemental information

Supplemental Figure 1-2 and Supplemental Table 1-3

## Author contributions

J.L., C.P., Y.T., Y.L. and Z.S.P., performed the experiments. M.E.S., Y.T., X.C., H.Y., and M.B. analyzed data. M.E.S. and M.B. wrote the manuscript. M.W. and M.B. provided supervision.

M.B. conceptualized the study. All authors edited the manuscript.

