## Supplemental Figures and Supplemental Table 1 for "Convergent DNA Methylation Abnormalities at Bivalent Chromatin in Human Growth Disorders"

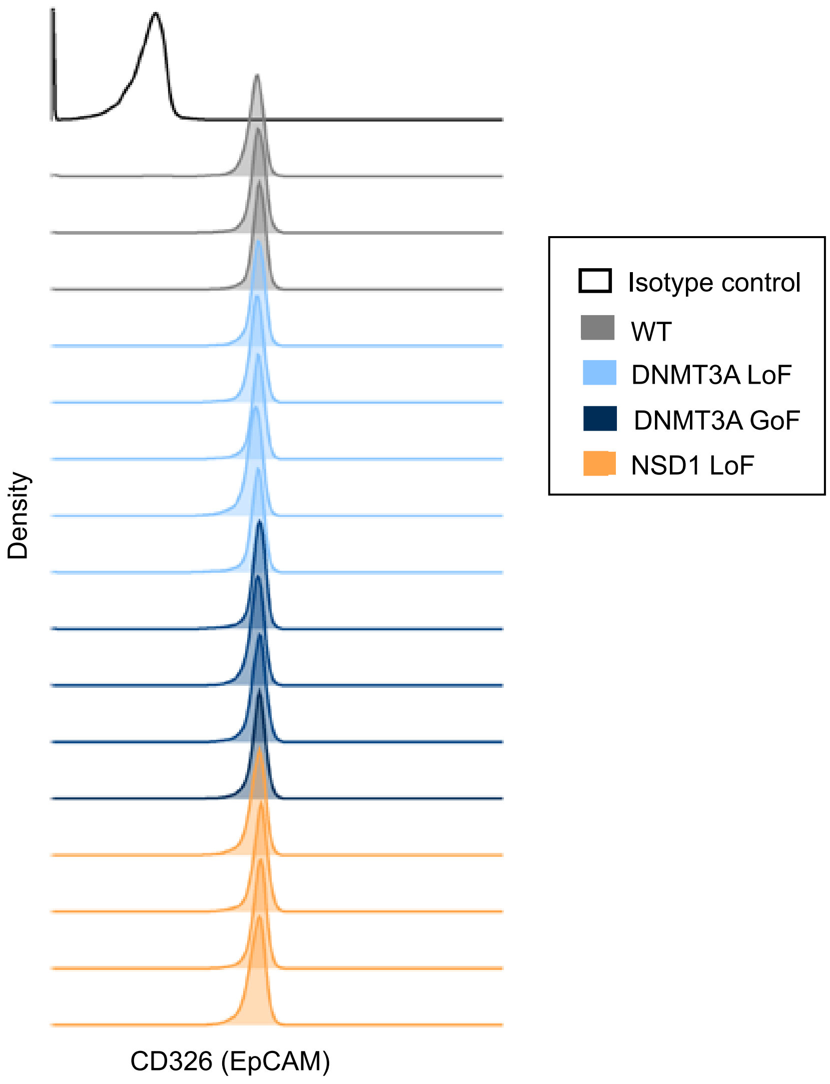


**Supplemental Figure 1. Validation of pluripotency of hESCs.**
Expression levels of CD326 (EpCAM), a pluripotency marker, in WT and mutant hESC clones were assessed by flow cytometry.

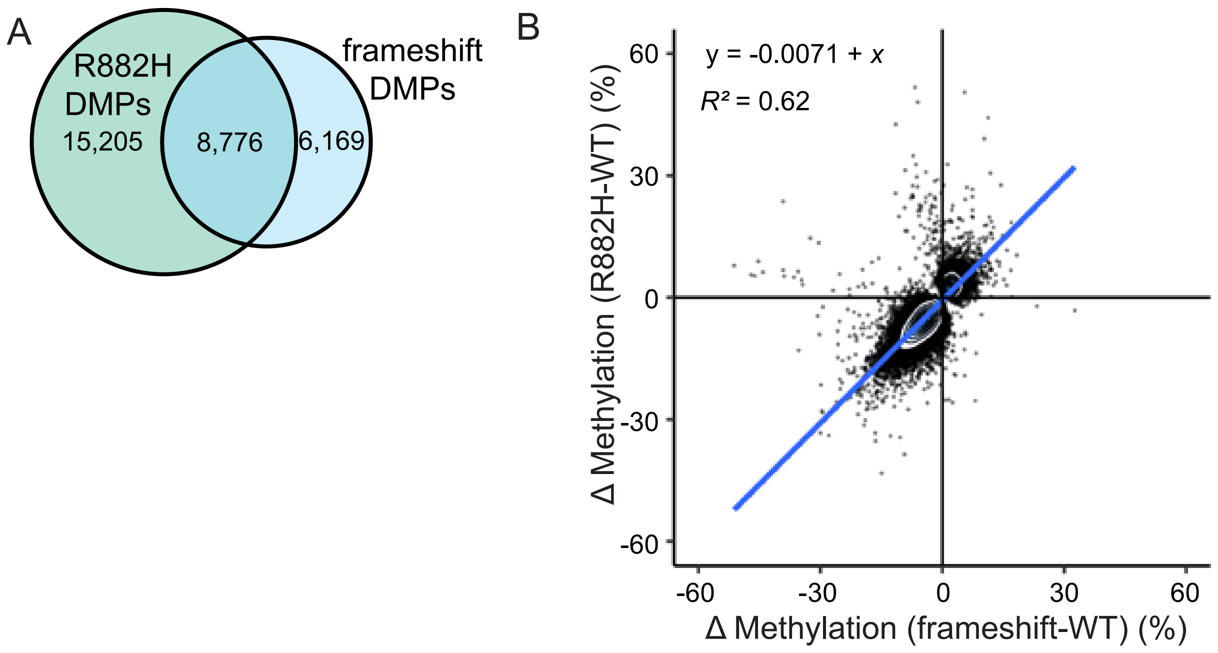


**Supplemental Figure 2. Comparison of differentially methylated positions (DMPs) in DNMT3A frameshift and R882H hESC clones.**

(A) Venn Diagram showing the overlap of DMPs between DNMT3A frameshift and R882H clones. P < 2.2*10-16, Fisher's exact test comparing frameshift-unique DMPs s to those shared with R882H among all probes.

(B) Pairwise comparison of methylation differences at all DMPs identified in DNMT3A frameshift and R882H clones relative to WT. Linear regression and Pearson's correlation coefficient ($R^2$) are shown.

| Syndrome | GEO accession | Platform | Samples analyzed |
|---|---|---|---|
| TBRS | GSE128801 | Infinium Human Methylation 450K BeadChip | GSM3685724, GSM3685725, GSM3685726, GSM3685727, GSM3685728, GSM3685729, GSM3685731 |
| HESJAS | GSE120428 | Infinium MethylationEPIC v1.0 BeadChip | GSM3400677, GSM3400678, GSM3400680 |
| SS | GSE191276 | Infinium MethylationEPIC v1.0 BeadChip | GSM5742874, GSM5742875, GSM5742876, GSM5742877, GSM5742878, GSM5742879, GSM5742880 |

**Supplemental Table 1**. **Growth syndrome patients whose data was reanalyzed for this study.**
